# Effect of PEI-coated MNPs on the Regulation of Cellular Focal Adhesions and Actin Stress Fibres

**DOI:** 10.1101/617480

**Authors:** Kaarjel K. Narayanasamy, Joshua C. Price, Marwan Merkhan, Ajile Elttayef, Jon Dobson, Neil D. Telling

## Abstract

The biocompatibility of surface coated/functionalised magnetic nanoparticles (MNPs) is key to their successful incorporation and application in biological systems. Polyethylene imine (PEI) -coated MNPs provide improved *in vitro* transfection efficiency compared to conventional chemical methods such as Lipofectamine and cationic polymers, and are also safer than viral transduction. Commercial cell toxicity assays are useful for end-point and high-throughput screening, providing fast results and an overview of cell health. However these assays only take into account cells that have undergone an extreme toxic response leading to cell death. Cell toxicity is a complex process which can be expressed in many forms, through morphological, metabolic, and epigenetic changes. A common indicator of cell stress and toxic response is increased cell adhesion and stress fibre formation. It is important to identify these changes in cells as it may affect downstream results and applications in biomedicine. This study explores the effect of the nanomagnetic transfection agent PEI-coated MNPs (MNP-PEIs) and an external magnetic field on cell behaviour, by studying particle internalization, changes in cellular morphology, and cell adhesion. We found that MNP-PEIs induced cell stress through a dose-dependent increase in cell adhesion via the overexpression of vinculin and formation of actin stress fibres. While the presence of PEI was the main contributor to increased cell stress, free PEI polyplexes induced higher toxicity compared to PEI bound to MNPs. MNPs without PEI coating however did not adversely affect cells suggesting a chemical effect instead of a mechanical one. In addition, genes identified as being associated with actin fibre regulation and cell adhesion, showed significant increases in expression due to the internalization of the MNP-PEI complex. From these results, we identify anomalous cell behaviour, morphology, and gene expression after interaction with MNP-PEIs, as well as a safe dosage to reduce acute cell toxicity.

## INTRODUCTION

The field of biomedical science makes use of magnetic nanoparticles for a large variety of applications. The ease of manipulation of MNPs make them particularly attractive for *in vivo* applications, as researchers continue to seek methods to improve diagnosis and treatment non-invasively. Applications include the use of MNPs as contrast agents for magnetic resonance imaging (MRI)^1,2^, for drug and gene delivery^3–5^, magnetic separation of cells^6^, and hyperthermia^7–9^. As such for MNPs to be suitable for biological applications and safe in a clinical setting, they should exhibit biocompatibility, low toxicity, and chemical stability.

Studies using standard cell endpoint assays have shown increased cytotoxicity of uncoated or bare MNPs^10^. Therefore, surface coating of MNPs are not only used to confer functionality, but to also to improve biocompatibility and protect cells from rapid biodegradation of MNPs and the production of ROS. For example, Baber et al. demonstrated that a silica shell around uncoated MNPs reduces the formation of soluble iron which improves cell viability ^11^. Although many surface coatings improve biocompatibility of MNPs, toxicity is also coating size and surface charge dependent ^10^.

MNPs are used commercially for *in vitro* applications such as transfection, cell labelling, cell separation, and magnetic 3D cell culture. Many companies selling MNPs for commercial purposes simply state that the particles are safe and elicit low to no cytotoxic effect on cells. This is usually demonstrated through standard cell viability or cell death assays, which typically conclude that cells respond well to MNP internalization. However, these assays do not take into account disruptions in cellular processes or downstream changes in cell behaviour which might affect the outcome of an experiment or application. Hence an understanding of these changes is especially important when considering potential therapeutic applications.

Hoskins et al. have stressed the importance of conducting the correct assay for cell viability and toxicity determination as some commercially available kits record inaccurate assay readings. This occurs due to interaction of the chemical with MNPs^12^, as MNPs are known to quench fluorescence signals, or bind to assay molecules. Although the mechanism of fluorescence quenching has not been elucidated, hypotheses behind the quenching may be due to a nonradiative energy transfer between the dye and MNPs through electron couplings, collisions between the dye and MNPs, or the broad absorption of fluorescence by iron oxide ^13,14^.

For applications in transfection, the rapid advancement of the CRISPR-Cas9 gene editing complex has seen breakthroughs in biomedical research, however its progress is limited to the effective but unsafe viral delivery methods. To this end, nanoparticles are being studied as a method to provide an alternative efficient and safe gene delivery system^15^. Nanomagnetic transfection is the non-viral delivery of genes into a cell using MNPs complexed with DNA and a magnetic field to attract the MNP-complex onto the surface of the cell^16,17^. Nanomagnetic transfection using an oscillating system was developed by the Dobson group, where the lateral movement of magnet arrays significantly improved transfection of cells in vitro^18–20^. PEI-coated MNPs (MNP-PEIs) are an effective gene delivery agent compared to conventional chemical transfection methods. They are also less toxic, have targeting capabilities and are multi-functional. However, in many cases MNPs do not elicit a simple live or dead toxicity result but instead cause underlying disruptions in cellular processes^21,22^. A study by Hoskins et al. studying the effect of MNP-PEIs on cell membrane integrity and cell stress based on endpoint assays^12^, however the phenotypic and genotypic changes are still unidentified as well as the components of MNP-PEIs responsible for cell stress. Furthermore, studies focused on commercial MNPs^22^ are limited in terms of understanding the cellular effect of individual components that make up the MNPs such as the protein corona, surface coating, and MNP cores.

By preparing and performing detailed characterization of the MNP-PEIs, we are able to assess the cellular changes in response to varying concentrations of MNP-PEIs and its individual components. To address the gap in toxicity studies for MNP-PEIs, this study seeks to determine the toxic response of cells when exposed to MNP-PEIs by measuring changes in cell adhesion through morphological and gene expression studies. An oscillating magnetic array used to enhance transfection was also studied to determine mechanical effects on cell adhesion and stress. Changes in cell adhesion can signify various functional and structural anomalies to the cell, as apart from adhesion, the cell cytoskeleton is also involved in structural support from external forces, transmission of external environmental cues to the cell and vice versa, formation of pseudopodia for movement, reorganization of organelles in the cell, endocytosis, and cell division.

In the work presented here, the vinculin molecule was studied to determine changes in cell adhesion of MNP-PEIs treated cells. Vinculin is associated with focal adhesion formation and the mechanism of vinculin association with actin and integrins as well as vinculin binding sites and active conformation has been widely reported ^23,24^. Vinculin and F-actin are commonly studied in cell adhesion research, as one factor which causes F-actin stiffness is its binding to vinculin which initiates actin bundling and cross-linking ^25^. To date, four main stress fibres have been identified, which are the dorsal and ventral stress fibres, transverse arcs, and the perinuclear actin ^26,27^.

We focus here on documenting changes in cellular morphology as well as gene expression profiles relating to changes in focal adhesion and F-actin of cells interacting with MNP-PEIs. The individual components of the MNP-PEI complex, i.e. free PEI molecules and oleic acid MNPs, are also separately studied to determine the aggravating factors contributing to cell stress. Besides chemical stimuli, mechanical/physical effects arising from the use of an external magnetic force on MNP-PEIs will also be discussed. Finally, the dose dependent effect of MNP-PEIs on cells is also addressed, to identify safe concentrations for *in vitro* biological applications.

## RESULTS

The preparation and characterization of MNP-PEIs used in this study are described in detail elsewhere^28^. Briefly, MNPs were prepared using the thermal decomposition method^4^ to obtain largely spherical monodisperse particles with a mean core size of 17.4 nm and 15% polydispersivity. These MNPs have a negatively charged oleic acid coating on their surface (oleic acid MNPs). The MNPs were then coated with polyethyleneimine (PEI) by ionic interaction to obtain a dense PEI surface coating, with high MNP colloid stability and a positive surface charge (Figure 1). The PEI surface coating confers functionality to MNPs, where it can be used for nanomagnetic transfection.

**Figure 1:**
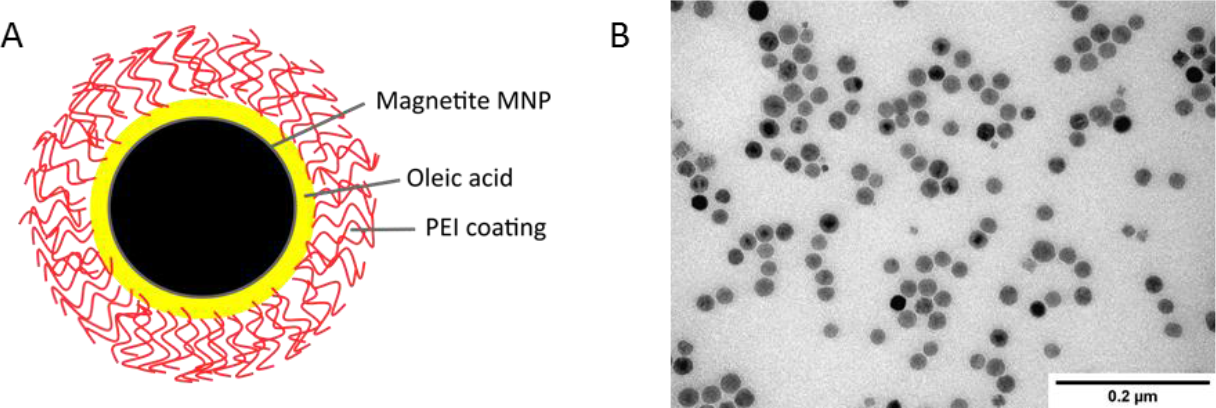
(A) Schematic of MNP-PEI consisting of a magnetite core with a negatively charged oleic acid shell and a surface coating of positively charged PEI polymer. (B) TEM image of MNPs with a largely spherical shape.

### Changes in cell density and morphology

Trypsin is a proteolytic enzyme which is used in cell culture to cleave adhesion proteins and detach adherent cells from the cell culture surface. Brightfield images of MG-63 cells treated with MNP-PEIs on oscillating magnet arrays are shown in Figure 2, before and after treatment with trypsin.

**Figure 2:**
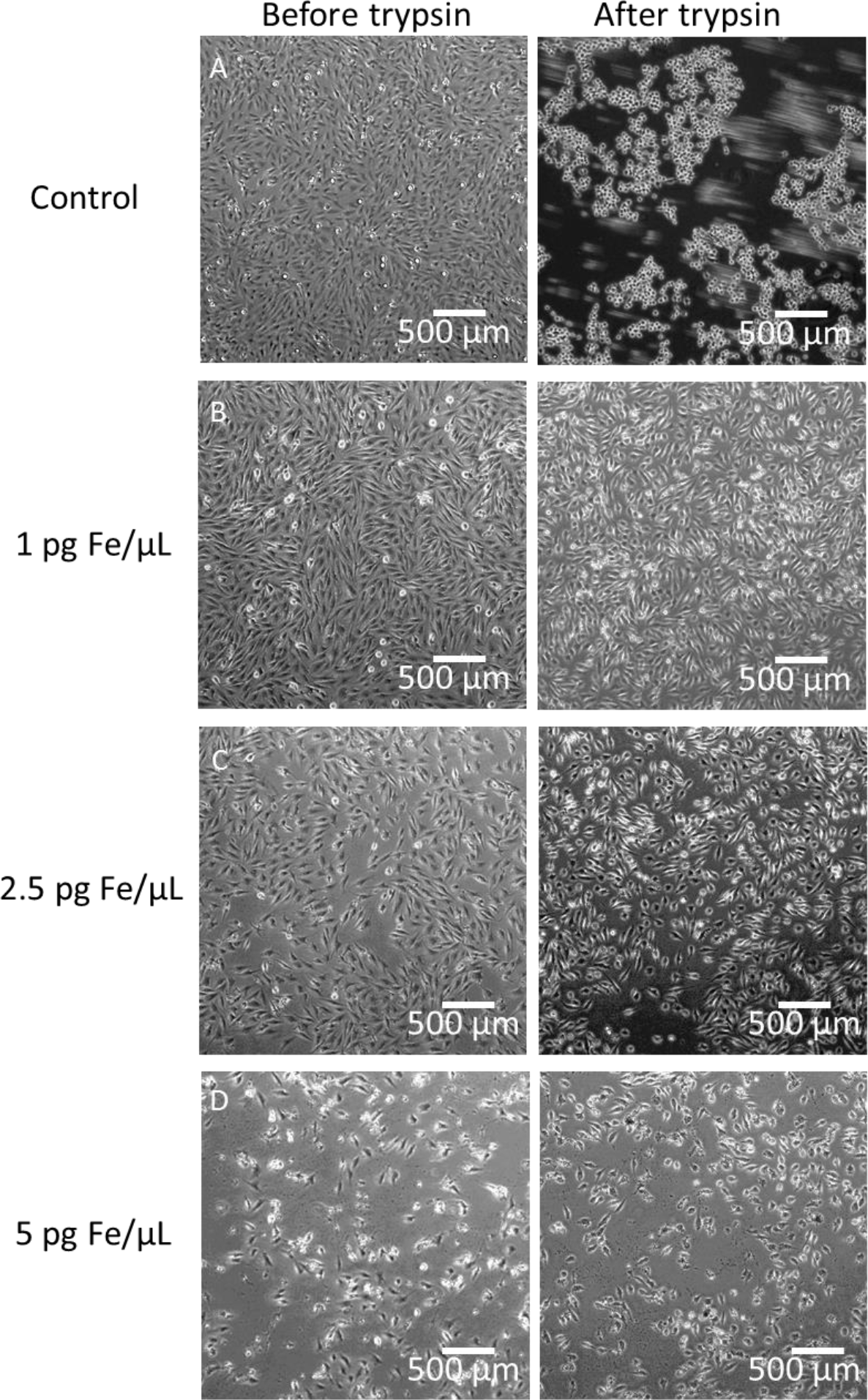
Brightfield images of MG-63 cells with MNP-PEI in T75 flasks before and after treatment with trypsin for cell detachment. Treatment groups are (A) control cells without MNP PEI, cells treated with (B) 1 pg Fe/μL, (C) 2.5 pg Fe/μL, and (D) 5 pg Fe/μL of MNP-PEI. Cells were completely detached in the control group but not in the flasks containing MNP-PEI.

In Figure 2A (left column), cells not treated with MNP-PEIs were confluent, with a typical spread morphology before the addition of trypsin. Incubation with trypsin completely detached cells (Figure 2A, right column) which can be observed from the spherical morphology of cells floating in the culture media. By contrast, whilst cells treated with 1 pg Fe/μL MNP-PEIs (Figure 2B) were also confluent and had a typical spread morphology, they did not show detachment after trypsin treatment. Similarly, the 2.5 and 5 pg Fe/μL MNP-PEI treatment groups showed little trypsin activity as most cells still adhered to the bottom of the flask (Figure 2C and D).

At a lower dosage of MNP-PEIs (Fig. 2B), cells maintained a typical spread morphology but with increasing MNP-PEI dosage (Fig. 2C and D), cells showed reduced confluency which suggested reduced proliferation, as well as a rounded morphology when compared with untreated controls (Figure 2A). Signs of cell stress at 5 pg Fe/μL MNP-PEI (Fig. 2D) was noticeable, and the high dose resulted in a loss of cells possibly due to PEI toxicity. However the cells that survived, adhered strongly to the flask.

To assess the internalization of different doses of MNP-PEIs in MG-63 cells on oscillating magnet arrays, the Prussian blue assay was employed to observe intracellular iron content (Figure 3A). The ferrozine assay was then used to quantify intracellular Fe^3+^ ions (Figure 3B), which saw a significant increase in MNPs internalized in cells with increasing MNP-PEI dosage of 0.14, 0.4, and 0.7 pg Fe/μL. Cells were treated with MNP-PEIs for 30 minutes and thoroughly washed to remove non-associated MNP-PEIs. Assays were performed after 24 hours.

**Figure 3:**
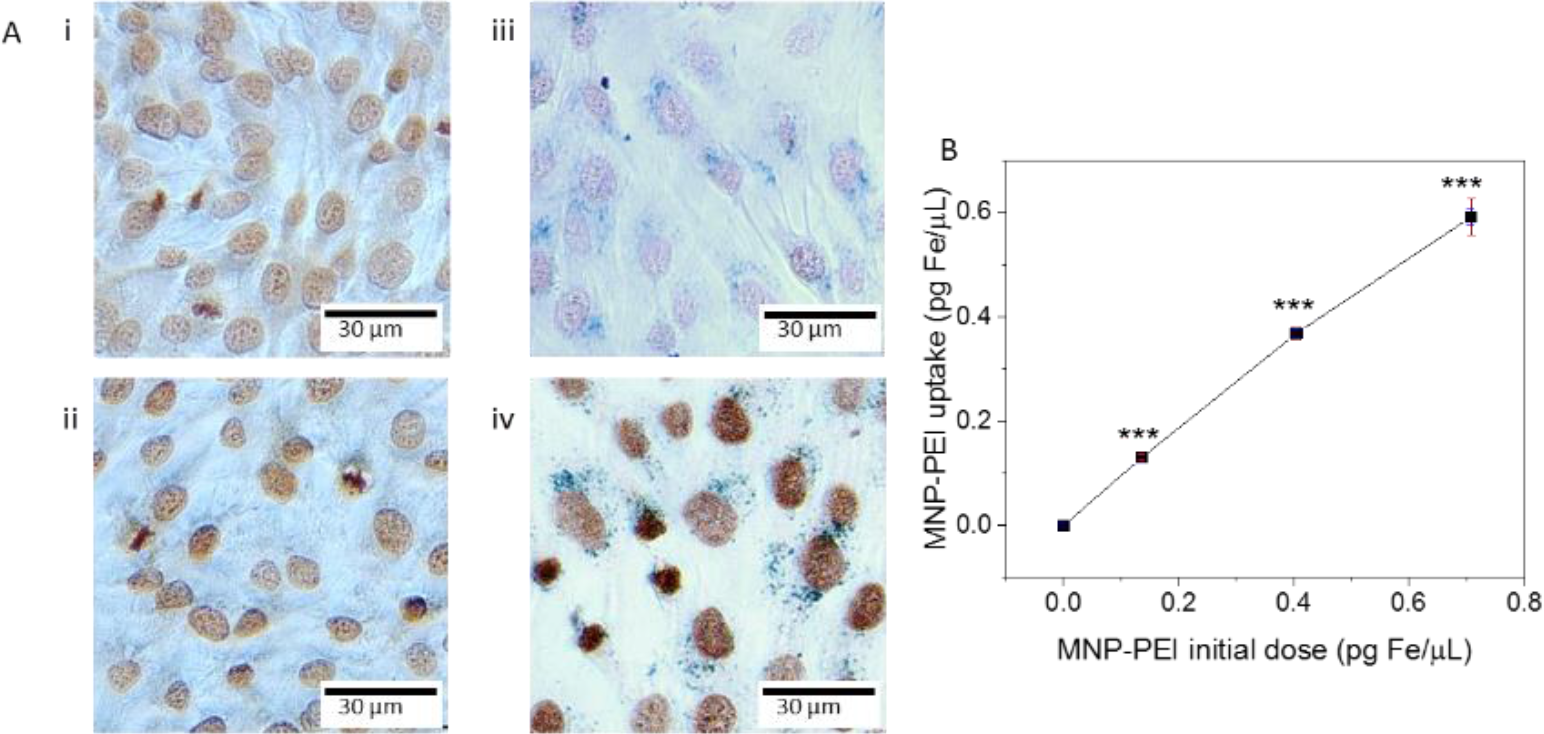
MG-63 cells treated with MNP-PEIs for 30 minutes, washed and incubated for 24 hours. (A) Brightfield images of cells with Prussian blue for iron staining in cells (blue) and counterstained with nuclear fast red. (i) Control, (ii) 0.14 pg Fe/μL, (iii) 0.4 pg Fe/μL, (iv) 0.7 pg Fe/μL of MNP-PEIs per well. (B) Quantification of MNP-PEI dose vs internalization in cells using ferrozine assay, N=3, ***p<0.0005.

The mass of MNP-PEIs taken up by cells appeared to be dose dependent. The control group at 0 pg Fe/μL showed negligible iron concentration detected in cells. At low concentration of MNP-PEIs (0.14 pg Fe/μL), particles were not visible in cells using Prussian blue staining. The ferrozine assay (Figure 3B), however, demonstrated that cells contained a small amount of iron after 24 hours of incubation. At a higher concentration of MNP-PEIs (0.4 pg Fe/μL, Fig. 3A-iii), the particles were discernible as fine blue particulate distributed within the cytoplasm. At the highest dosage (0.7 pg Fe/μL, Fig. 3A-iv), the particles were visible as large aggregates, indicated by granular regions in cells which clustered around the perinuclear region.

Ferrozine measurements of the iron content in the cells revealed that the uptake of particles increased significantly (p=0.0028) with each MNP-PEI dosage. As expected, the amount of MNP-PEIs taken up by cells were directly proportional to the initial dosage of MNP-PEIs. This effect however was finite at an MNP-PEI concentration of 0.7 pg Fe/μL as there was a significant difference between particles seeded in cells and the lower concentration taken up by cells (p=0.0004).

Uptake of particles into cells was close to saturation in the 0.7 pg Fe/μL MNP-PEI group. One limiting factor to particle uptake is the rate of endocytosis, therefore with a longer incubation time and a higher concentration of MNP-PEIs, particle uptake would increase but at the cost of cell health. This effect has been described previously as overstimulated endocytosis^21^ which implies a positive feedback effect of MNP dosage versus rate of endocytosis. MNP-PEIs are also toxic to cells at high concentrations, therefore cells can only take up a low dose before undergoing cell death. Furthermore, cells treated with higher doses of MNP-PEIs were observed to have lower cell density, related to reduced proliferation (Figure 3A).

### Components of MNP-PEI inducing cell adhesion

A PicoGreen DNA quantification assay was used to investigate cell adhesion by quantifying the number of cells which remain attached in cell culture wells after treatment with trypsin and removal of detached cells. The amount of DNA represents the population of live cells unaffected by trypsinization, following removal of unbound cells. Therefore, higher PicoGreen fluorescence intensity indicated stronger cell adhesion.

Figure 4A compares cell adhesion on two cell types, which are MG-63 and HeLa cells after MNP-PEI treatment. The effect on cell adhesion of the components of the MNP-PEI complexes, i.e. oleic acid MNPs and free PEI was also studied. Figure 4B shows MG-63 cells treated with oleic acid MNPs and Figure 4C are cells treated with PEI polyplexes. The treated cells were either assayed immediately after 30 minutes on the magnet array, or after 30 minutes on the magnet array and 24 hours of incubation.

**Figure 4:**
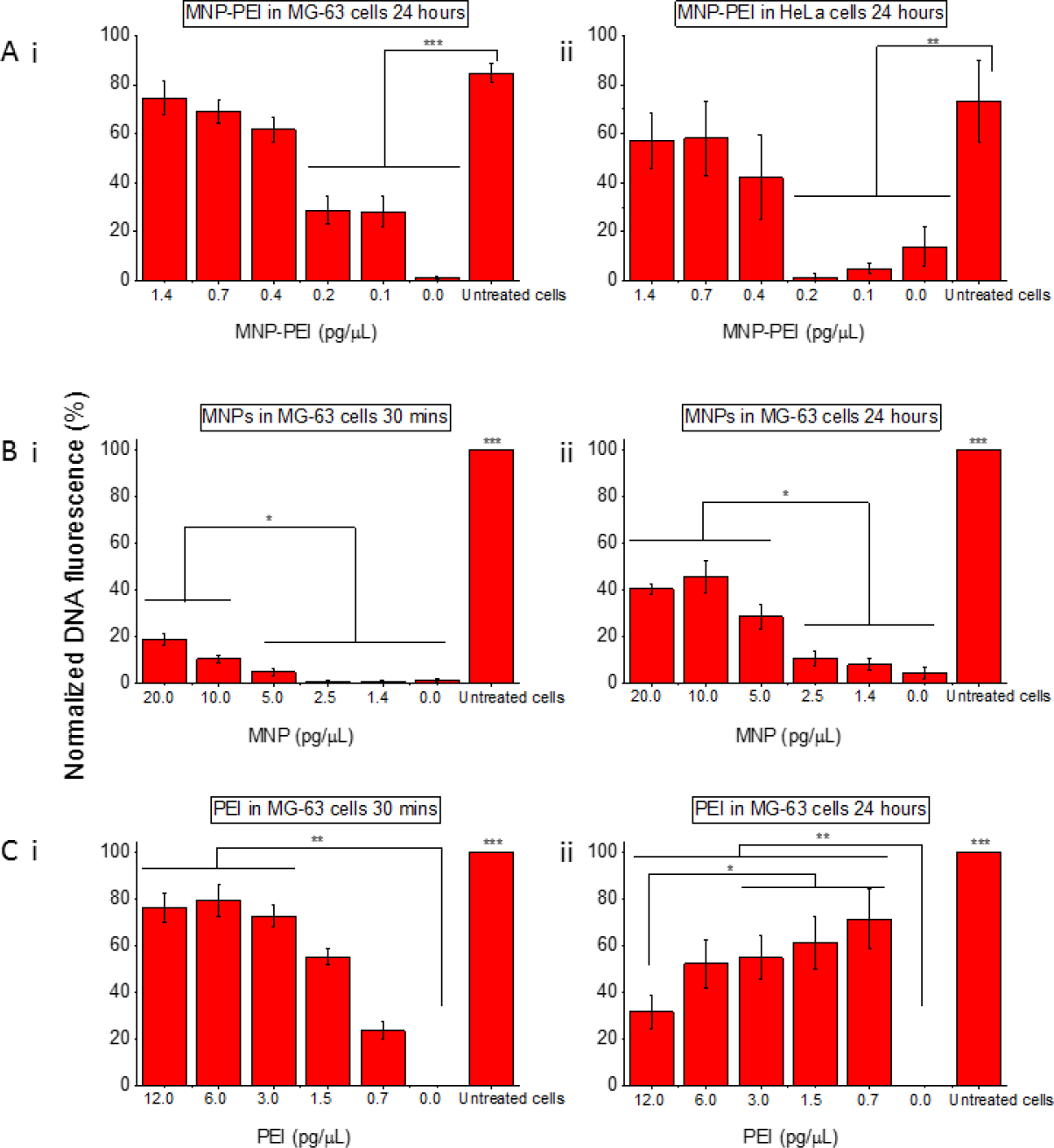
Pico green assay to determine the relative number of cells adhered to cell culture wells after cells were loaded with MNP-PEIs, oleic acid MNPs, or PEI only, and subsequently treated with trypsin to detach cells. (A) (i) MG-63 cells and (ii) HeLa cells seeded with MNP-PEIs on a magnet array for 30 minutes and 24 hour incubation. (B) MG-63 cells with oleic acid MNPs 30 minutes on a magnet array (ii) and 24 hour incubation. (C) MG-63 cells with PEI polymer 30 minutes on the magnet array (ii) and 24 hour incubation. N=5, *p<0.05, **p<0.005, ***p<0.0005.

#### Effect of MNP-PEI dosage on two cell types

HeLa cells were compared with MG-63 cells to determine if the adhesion effect persisted in more than one cell type. Figure 4A-i and ii compares the adhesion between the two distinct cell types, when cells were detached with trypsin treatment after 24 hours incubation with MNP-PEIs. Both cell types showed the same effect when treated with MNP-PEIs at high dosage, where cells treated with 1.4, 0.7, and 0.4 pg Fe/μL MNP-PEI had high adhesion to the cell culture surface, whereas lower doses of ≤ 0.2 MNP-PEI pg Fe/μL per well showed detachment of cells after trypsinization (MG-63 cells p< 1.3e^−6^, HeLa cells p< 0.006).

Although both cells showed a change in adhesion from MNP-PEI interaction, HeLa cells exhibited lower cell adhesion compared to MG-63 due to the ability of osteoblasts to deposit collagen for increased binding to the extracellular matrix (ECM).

#### Effect of incubation time

For cells treated with MNPs in the ‘30-minute incubation time’ group (Fig. 4B-i), changes in cell adhesion were small. However, high MNP dosage (10 and 20 pg Fe/μL) showed significant differences to the other treatment groups (p< 0.04). When MNPs were incubated with the cells for 24 hours (Fig. 4B-ii), there was a significant increase in cell adhesion for the 3 treatment groups with the highest seeded MNP doses (p< 7.6e^−4^).

PEI after 30 minutes (Fig. 4C-i) increased cell adhesion for high PEI doses (12, 6, and 3 pg/μL; p< 9.5e^−7^). Interestingly, cells treated with PEI for 24 hours at the highest PEI concentration of 12 pg/μL had significantly lower amounts of DNA (p< 0.034) (Fig. 4C-ii) due to increased cell death and detachment. This indicates a cytotoxic effect of PEI at higher dosage with long incubation time. Low dosage of PEI (1.5 and 0.7 pg/μL) incubated in the cell culture for 24 hours had a stronger influence on cell adhesion compared to the 30-minute treatment.

Therefore, oleic acid MNPs had a minimal effect on cell adhesion, whereas cell adhesion increased almost immediately when exposed to PEI polyplexes. PEI also caused cell death when in contact with cells for a prolonged period of time.

#### Effect of MNPs or PEI polymer

The effect of the individual components of MNP-PEIs on increased cell adhesion was demonstrated in Figure 4B and 4C. While cells displayed increased adhesion with a minimum of 0.4 pg Fe/μL MNP-PEI treatment, the same effect was not observed when cells were treated with oleic acid MNPs at the same concentration. Cells treated with PEI, however, had an adverse effect, where low concentrations of PEI were able to cause a strong cell adhesion response in a short time.

The lowest concentration of MNP-PEIs that induced cell adhesion was at 0.4 pg Fe/μL MNP-PEI which has a surface coating of approximately 4 pg/μL PEI. However cells incubated with free PEI had an increase in adhesion with only a fifth of the PEI concentration (0.7 pg/μL PEI).

Therefore, the key factor of MNP-PEI induced cell adhesion is the PEI surface coating. Interestingly, bound PEI is shown to have reduced cytotoxicity compared to free PEI polyplexes in the cell at the same dosage.

### Effect of MNP-PEI dosage on vinculin and actin stress fibre formation

MNP-PEIs can be used as transfection agents, where the binding of DNA to the surface of MNP-PEI enables transfection. MG-63 cells were transfected using a GFP reporter plasmid to observe successful transfection. The study below was conducted to assess uptake and stress response of cells to the different components of the MNP-PEI transfection agent.

The difference in particle type taken up by cells is noticeable in brightfield images in Figure 5. In cells treated with MNP-PEIs and MNP-PEIs carrying DNA (Fig.5-iii&iv), dark granules are observed in the cytoplasm and perinuclear region of cells, indicating uptake of MNPs into cells. Cells treated with oleic acid MNPs (Fig. 5-ii), however lack this structure. Furthermore, the formation of filopodia can be seen in cells treated with MNP-PEI and MNP-PEI carrying DNA.

**Figure 5:**
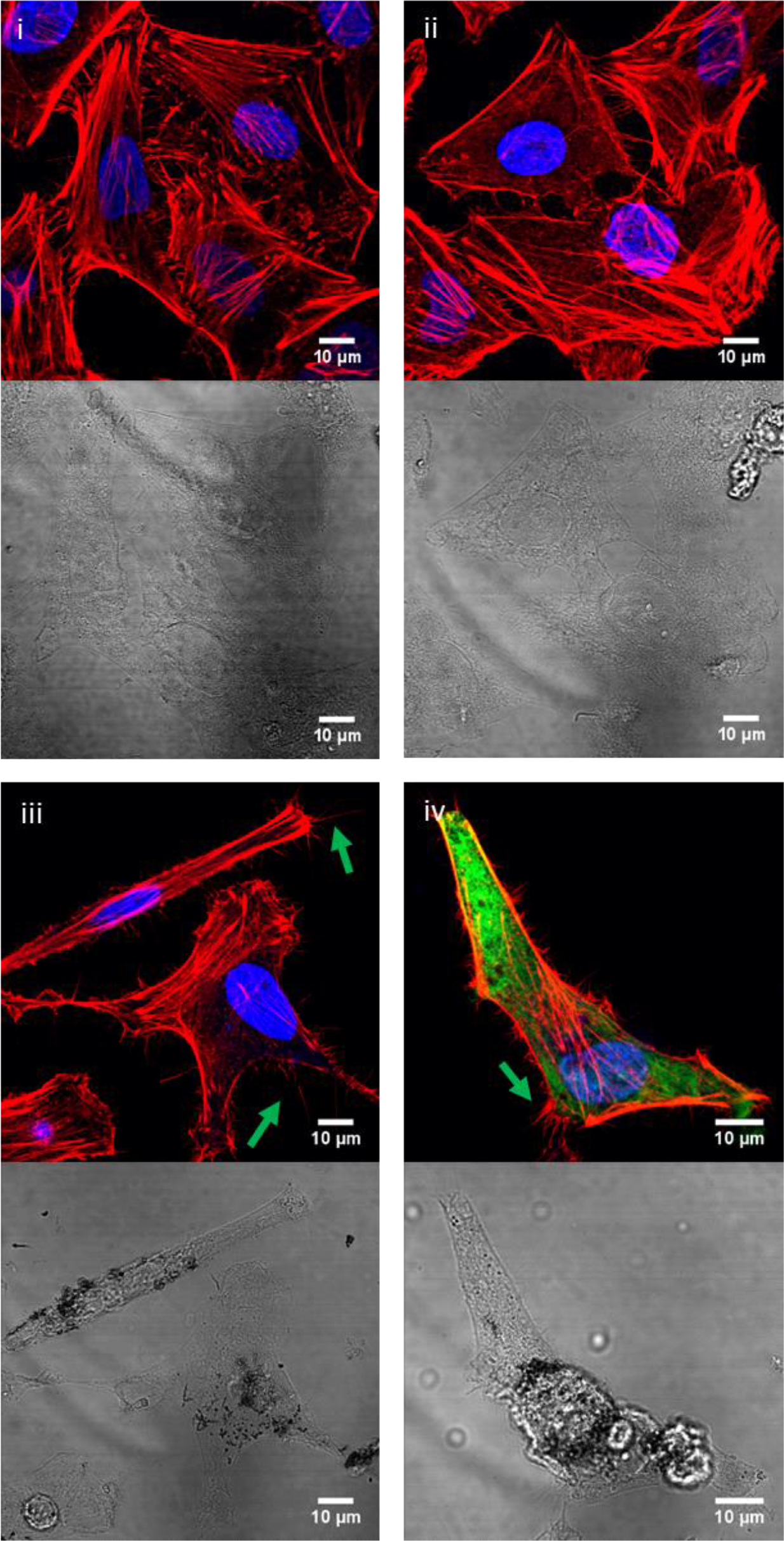
Fluorescence (top) and brightfield images (bottom) of MG-63 cells (i) control, or treated with (ii) oleic acid MNPs, (iii) MNP-PEIs, and (iv) MNP-PEIs with GFP plasmid. Cells are stained with actin (red) and nucleus (blue), and green fluorescence (in Fig. C iv) shows the successful transfection of cells with the GFP plasmid. Green arrows indicate filopodia.

The difference in uptake and changes in morphology can be attributed to the presence of PEI on the surface of MNPs. Oleic acid MNPs without PEI coating carry a negative charge, and have a relatively weaker interaction to cells compared to PEI-coated MNPs which carry a high positive charge density^10,30^, hence oleic acid MNPs did not induce strong cell adhesion. The MNP-PEI complex is able to transfect cells successfully (Figure 5-iv), however the interaction of cells with MNP-PEIs caused cell stress as evidenced by the protrusion of filopodia and the unusual morphology of the cell in the brightfield image. This dose dependent effect is studied further below.

The effect of MNP-PEIs on cell adhesion was assessed using immunostaining for vinculin. Additionally, phalloidin based F-actin staining was employed to assess actin stress fibre formation. A striking increase in vinculin intensity was observed for cells treated with higher doses of MNP-PEIs, whereas cells incubated with lower doses (0.14 pg Fe/μL MNP-PEIs) had a similar staining profile to control cells (Figure 6A). Increasing doses of MNP-PEIs (0.4 and 0.7 pg Fe/μL MNP-PEIs) significantly increased vinculin fluorescence intensity compared to control cells (p=6.4e^−7^) and vinculin was localised predominantly in the nucleus but also expressed in the cytoplasm. Fluorescence imaging of F-actin (Figure 6A, right column) showed a comparable difference between the control and 0.14 pg Fe/μL groups (Figure 4A i and ii) with the 0.4 pg Fe/μL and 0.7 pg Fe/μL groups (Figure 6A iii and iv), where randomly oriented actin cytoskeleton preceded a reorganization of the actin cytoskeleton to form ordered parallel stress fibres. Dendritic protrusions of F-actin from the cell membrane (green arrows) was observed in MNP-PEI groups.

**Figure 6:**
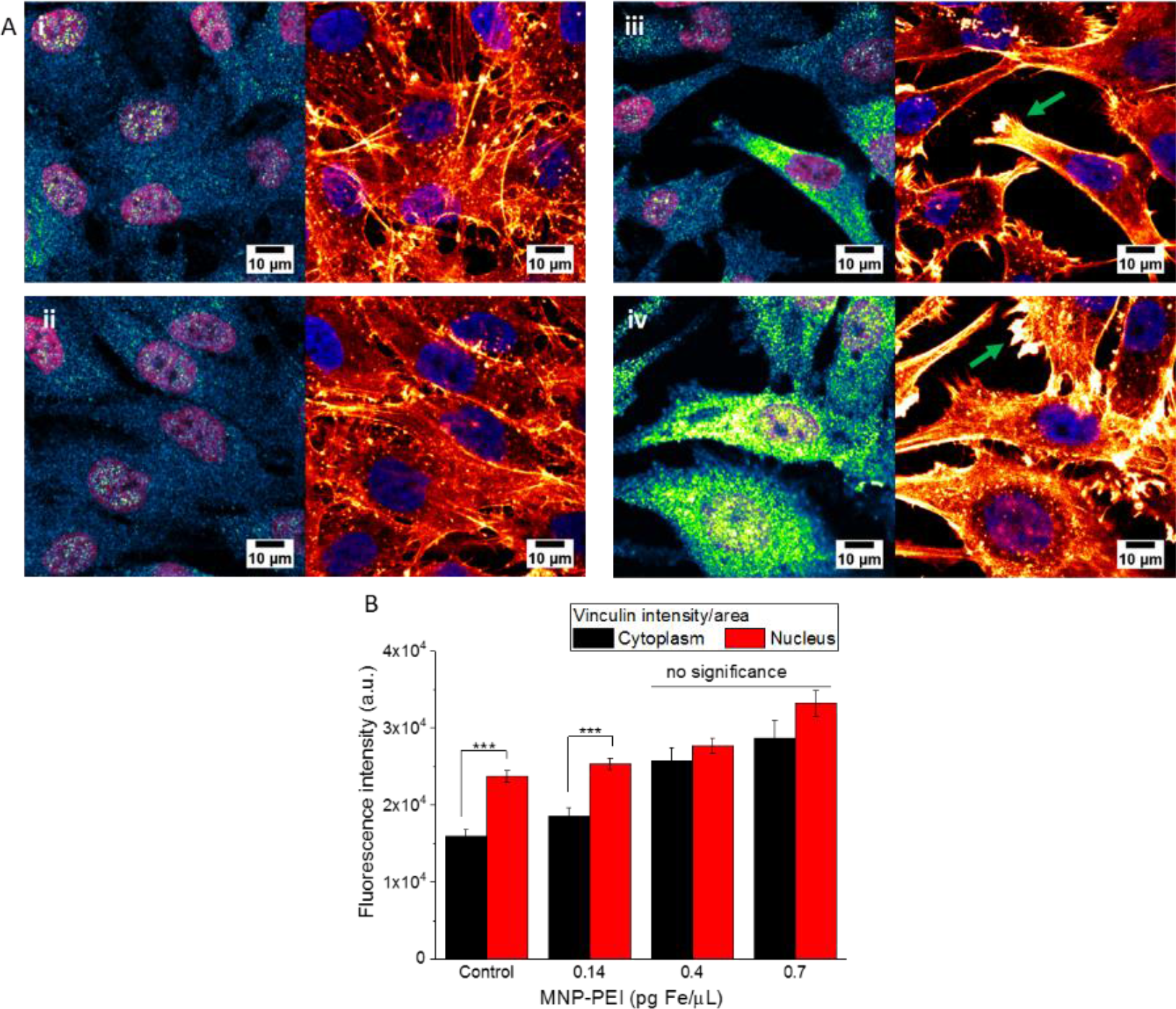
(A) Fluorescence images of nucleus (purple), vinculin in blue-green (left), and F-actin in red (right). MG-63 cells were the untreated (i) control, or treated with (ii) 0.14 pg Fe/μL MNP-PEI, (iii) 0.4 pg Fe/μL MNP-PEI and (iv) 0.7 pg Fe/μL MNP-PEI. The fluorescence change from blue to green to yellow indicates increasing vinculin concentration. (B) Quantification of vinculin fluorescence intensity per area in the nucleus and cytoplasm showing significantly higher vinculin expression at high MNP-PEI doses per well (0.4 and 0.7 pg Fe/μL) N = 25 to 40 cells.

Quantification of fluorescence intensity of vinculin in cells was used to compare between the cytoplasmic area and nuclear area to determine whether vinculin formation during cell stress occurs randomly within the cell or is localized at the centre or periphery of the cell. Overall, the expression of vinculin was higher in the nuclear region of cells compared to the cytoplasm in all groups (Figure 6B). A significant increase in vinculin expression was observed in the cytoplasm with higher dosage of MNP-PEIs in cells (0.4 and 0.7 pg Fe/μL) (p< 0.001) compared to the control and 0.14 pg Fe/μL MNP-PEI groups, however vinculin expression in the nuclear region remained relatively unchanged until treated with 0.7 pg Fe/μL MNP-PEI (p< 0.005).

The difference in vinculin expression between the cytoplasmic and nuclear area per treatment group was only significant in the control and 0.14 pg Fe/μL groups (p< 0.005), and at higher MNP-PEI dosage the intensity of vinculin expression increased in the nuclear area and was comparable to cytoplasmic intensity. Overall, treatment of MNP-PEI 0.7 pg Fe/μL showed the highest expression of vinculin in cells, followed by 0.4 pg Fe/μL MNP-PEI. Analysis via two-way ANOVA showed that the population means for each variable was significantly different (MNP-PEI dosage, p=5e^−15^, cytoplasm to nuclear area intensity, p=2.6e^−8^), but MNP-PEI dosage in cells is not significant to the localization of vinculin. The increase in vinculin intensity surrounding the nuclear area may be due to a specific set of stress fibres, known as the perinuclear actin cap. These stress fibres attach to focal adhesions and form around the nuclear region to maintain the shape and position of the nucleus in the cell, as well as transmit forces from the ECM to the nucleus ^32,33^.

### Gene expression profile

To further elucidate the effect of MNP-PEIs on cell adhesion and stress, gene expression analysis by qPCR was performed on cells treated with a low dose (0.5 pg Fe/μL) and high dose (1.0 pg Fe/μL) of MNP-PEIs. Seventeen genes were identified as those associated with the cell adhesion pathway and actin fibre remodelling and were selected for qPCR analysis.

**Figure 7:**
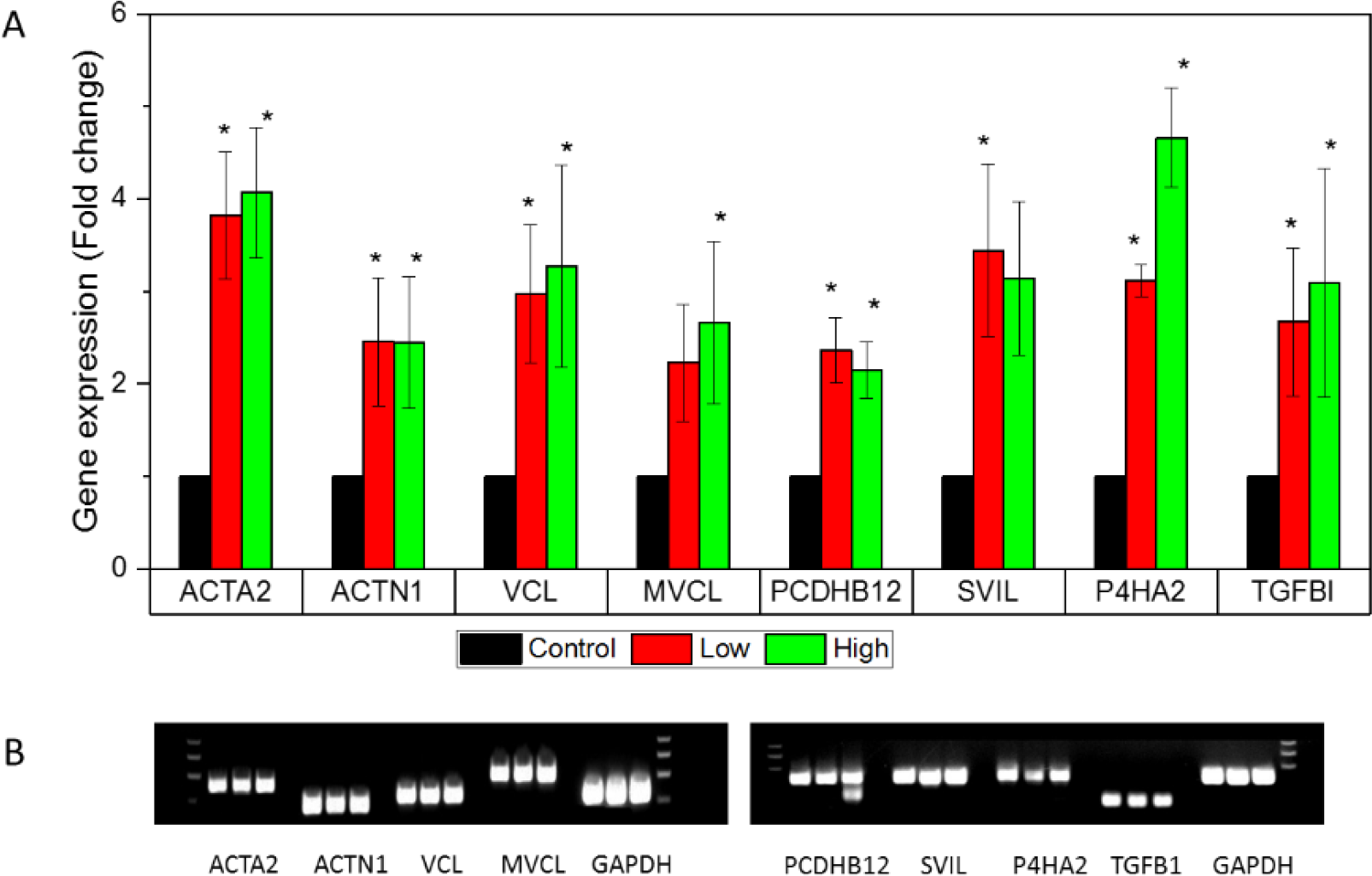
(A) Quantitative PCR of genes involved in cell adhesion and the cytoskeleton showing the effect of MNP-PEIs in cells. Treatment of MNP-PEIs in low dose (0.5 pg Fe/μL) (green) and high dose (1.0 pg Fe/μL) (red) is compared to control cells untreated with MNP-PEIs (black) and expressed in fold changes. F tests and student's t-tests were performed using ΔCt values, N=3-4, *p<0.05. (i) Genes ACTA2, ACTN1, MVCL and VCL and (ii) P4HA2, PCDHB12, SVIL and TGFB1. (B) Gel electrophoresis of qPCR products indicating the purity of qPCR products with each band group showing control (left), low MNP-PEI (middle), and high MNP-PEI (right). GAPDH was used as a housekeeping gene.

qPCR data of gene expression for the selected genes (Figure 7A) show upregulation in cells treated with MNP-PEIs. Out of the 17 genes, 8 were significantly upregulated compared to untreated controls (p<0.05). Interestingly, particle dosage (low and high) did not have a significant difference in gene upregulation. Compared to the untreated controls, treatment with MNP-PEIs resulted in an upregulation of all genes by a fold-change of more than 2.

To determine the specificity of the primers used for qPCR a melt curve was obtained from the qPCR instrument. The qPCR products were then run on agarose gel electrophoresis. GAPDH was used as a housekeeping gene. A primer that binds and amplifies a specific product has only a single peak in the melt curve, as well as a single band in the agarose gel electrophoresis. Double DNA bands indicate non-specific binding of primer and amplification of sample. Based on Figure 7B, all the products of qPCR reactions showed a single band except for gene PCDHB12 high dose.

## DISCUSSION

We have shown that there is a dose-dependent increase in cell attachment to their substrate in the presence of MNP-PEIs. This effect is caused by the PEI surface coating, while the MNP cores with an anionic oleic acid coating do not induce a cellular reaction at the same concentration. Increased focal adhesion complexes and rearrangement of the actin cytoskeleton was observed, as well as reduced proliferation and the formation of filopodia, indicating cell stress.

### Effect of MNP-PEIs on cell adhesion

In this study, a positive correlation was observed with increased MNP-PEI dosage in cells to the increase in vinculin expression and intensity of actin stress fibres. The mechanism with which cells increase their attachment onto a matrix is through the formation of focal adhesions. Focal adhesions are sites where the cell is connected to the external microenvironment through the anchoring of adhesion-associated proteins such as vinculin to cytoskeletal actin fibres. Besides focal adhesion localization, vinculin are also present in adherens junctions to mediate cell-cell adhesions. Another function of vinculin is to bind and bundle F-actin for the transduction of external stimuli ^34^, observed in the form of stress fibres.

The two components of the transfection agent (MNPs and PEI) were studied individually to determine the underlying cause of cell adhesion. MG-63 cells treated with either cationic PEI polymer or anionic oleic acid MNPs expressed different intensities of cell adhesion in response to each component. The effect of PEI was stronger and more damaging to cells even at PEI doses that were lower than MNP doses in cells.

Interestingly, free PEI induces a stronger response from cells compared to bound PEI at the same concentrations. Free PEI has a tendency to aggregate to form polyplexes, which has a higher positive charge density compared to bound PEI distributed over the large surface area of the MNPs. The high charge density causes higher acute toxicity, whereas PEI bound to MNPs may dissociate over time in the cell and therefore contribute to delayed toxicity^29^. Therefore, to improve overall cell health in biomedical applications, cationic polyplexes should be dispersed over a larger area to reduce their charge density.

The different response between cationic and anionic MNPs is due to the surface charge of the components, where positive charged surfaces have been shown to confer higher toxicity to cells^10^. PEI toxicity has been attributed to its destabilizing effects on the cell plasma membrane^35^. At lower concentrations however, PEI may induce the formation of focal adhesions and stress fibres (Fig 4c). The increased adhesion may be useful in some applications, where PEI can be used as a surface coating agent to enhance cell adhesion onto substrates such as higher attachment of some cell lines compared to collagen and poly-lysine^36^. For gene therapy and drug delivery applications, however, changes in cell morphology and gene regulation are not desirable effects, and maintaining the native state of a cell is preferable.

Apart from the biological incompatibility of PEI, endosomal lysis caused by PEI in cells^37^ also causes leaking of the acidic pH of endolysosomes into the cell cytoplasm, which could create an adverse reaction and toxicity in cells.

Adherent cells, unlike cells in suspension, need to attach to a surface before they regain normal function and remain viable ^38^. The effect of cell adhesion may be cell-type dependent as MG-63 cells showed noticeably stronger adhesion compared to HeLa cells. Osteosarcoma cells such as MG-63s are known to synthesize high amounts of various collagen types^39,40^. Cells have been documented to attach selectively to different collagen types^41^, and collagen is commonly used to improve adhesion of cells to tissue culture plastic^42^. The upregulation of prolyl-4-hydroxylase (P4HA) induced by MNP-PEIs indicates increased collagen synthesis by cells, and with the help of TGFBI, cells are able to anchor to extracellular collagen to strengthen cell adhesion to the ECM. Therefore MG-63 cells show more pronounced cell adhesion compared to HeLa cells when treated with MNP-PEI from the ability to synthesize and deposit collagen in the ECM.

### Effect of MNP-PEIs on the cytoskeleton

The actin cytoskeleton functions in various cellular processes such as cell spreading, migration, morphogenesis, cell division, and endocytosis ^33,43^. The uptake of material from the extracellular environment into the cell requires plasma membrane deformation and invagination to form endocytic vesicles. Therefore, one possible reason for the increased F-actin tensioning and increase in ACTA gene expression is to facilitate the capture of MNP-PEIs by endocytosis, hence inducing the formation of actin bundles with increasing MNP-PEI dosage.

The cytoskeleton is also involved in cell anchorage to the ECM by forming stress fibres with the help of focal adhesions. Stress fibres form by the polymerization of 10 to 30 actin fibres and myosin-II filaments by the gene alpha-actinin (ACTN), which was found to be upregulated with MNP-PEI treatment in this study.

Cellular protrusions known as filopodia were observed in cells treated with MNP-PEIs. Filopodia are formed by the bundling of actin fibres and function in cell-cell adhesion and motility, as well as for probing the extracellular environment^44^. Therefore, it seems likely that the pseudopodal extensions that were formed here, were in response to cellular stress to allow movement to suitable areas away from the foreign material causing the stress^45^.

Cell proliferation was observed to be affected by the presence of MNP-PEIs. This effect is evidence that MNP-PEI internalization interferes with actin-mediated regulatory processes as the actin cytoskeleton is also involved in processes such as proliferation and differentiation ^22^. This effect had been noted in previous studies with different cell lines, where iron oxide MNPs and polystyrene particles reduce proliferation ^46,47^ by interfering with cell metabolism and signalling ^48^.

### Chemical and mechanical stimuli

The stimulus that cells react to can either be chemical, though the interaction of PEI and MNPs, or through mechanical cues. Mechanotransduction has been widely reported to induce the formation of focal adhesions and cytoskeleton rearrangement^49,50^. Focal adhesions can form when mechanical stimuli from inside or outside the cell is present, which are transmitted through ion-channels or receptors located on the plasma membrane, thus recruiting adhesion molecules to the site for focal adhesion formation^51^. The ECM also receives mechanical forces from the external environment and transmits them to the cell through receptors and the cytoskeleton^52^.

Therefore besides chemical stimuli, mechanotransduction cannot be ruled out as the effect of an oscillating external magnet on MNPs in cells creates a downward force and lateral movement on the cell surface. Zheng et. al demonstrated a negative effect of mechanical forces on the proliferation and migration of cells ^53^. Previous studies report the manipulation of MNPs using external magnetic fields to exert rotating-oscillating motions or a simple magnetic pull downwards with a static magnet^54,55^ which caused cellular mechanotrasduction. In some cases, the development of focal adhesions to increase cell adhesion is desirable, where mechanotransduction using MNPs tethered to mesenchymal stem cells (MSCs) improved adhesion to the matrix as well as subsequent MSC differentiation^56^. The cell surface receptor that responds to MNP-magnetic forces was identified, which is the N-type mechano-sensitive calcium ion channel^57^. Although receptor-targeted magnetic mechanotransduction has applications in tissue engineering and regenerative medicine^58^, not much is known about chemical driven focal adhesion signalling caused by MNPs.

In this study, the oscillatory motion of MNPs on the surface of cells created by the oscillating magnet system and even the downward-directed force of MNPs on cells created using a static magnet^59^ may initiate mechanical cues for cells to recruit focal adhesion molecules. However the mechanical effect is obscured by the chemical stimuli exerted by PEI, since PEI without external magnets induced stronger attachment in cells compared to a high concentration of MNPs in the presence of oscillating magnets. For a significant mechanical effect to be observed, a higher concentration of MNPs and longer external magnet exposure may be required.

## CONCLUSION

Many studies have documented the toxicity profile of iron oxide NPs, however the particle coating is a significant contributing factor to cytotoxicity. This study demonstrates the increase in focal adhesion assembly and the formation of actin stress fibres in response to external chemical cues, i.e. MNP-PEIs. It also highlights higher PEI cytotoxicity as a free polymer compared to its bound state with MNPs, suggesting MNP-PEIs are a safer and more efficient transfection agent^60,61^ compared to PEI alone.

The effect of mechanical forces and substrate stiffness has been the main focus for the formation of stress fibres and focal adhesions^27^. This study provides an additional dimension to the mechanics of focal adhesion and stress fibre formation, where a similar cellular effect is observed arising from chemical cues from the external and internal environment.

For cell therapy applications, changes in cell signalling and processes should be studied thoroughly in addition to cell viability and toxicity assays. Future work should study the long-term implications of stress fibres and focal adhesion formation on physiological effects during nanomagnetic transfection.

## METHODS

### Cell culture

MG-63 and HeLa cells were cultured in Dulbecco’s Modified Eagle Medium (DMEM; high glucose with 4500 mg/L glucose, L-glutamine, sodium pyruvate, and sodium bicarbonate; Sigma) with supplemented 10% fetal bovine serum (FBS; Lonza) and 1% penicillin-streptomycin (10 000 units/mL; Lonza). Cells were seeded at a density of 35 000 cells/cm^2^ and grown to a confluency of 70 – 80% in 24 hours for treatments or passaged every 48 hours.

The MNP-PEIs used were synthesized using thermal decomposition, coated with PEI and characterized as described previously^28^.

### MNP-PEI treatment on MG-63 or HeLa cells

A stock suspension of MNP-PEIs (0.35 μg Fe/μL) was added into cell cultures at concentrations of 1, 2.5, and 5 pg Fe/μL respectively and placed on a horizontal oscillating magnet array (magnefect-nano from nanoTherics Ltd) at a frequency 2 Hz and 0.2 mm amplitude for 30 minutes. The media was replaced with fresh media. Cells were incubated for 24 hours until treated with trypsin and imaged.

In 24-well plates, either MG-63 or HeLa cells were treated with media supplemented with 0.14, 0.4, or 0.7 pg Fe/μL MNP-PEI, placed on an oscillating magnet array at a frequency 2 Hz and 0.2 mm amplitude for 30 minutes and incubated for 24 hours prior to assays. For gene expression studies, MG-63 cells were treated with 0.5 pg Fe/μL (low dose) and 1.0 pg Fe/μL (high dose) of MNP-PEI. RNA extraction was performed 24 hours after MNP-PEI treatment.

### Immunocytochemistry (ICC) for vinculin

MG-63 cells were cultured on glass coverslips and fixed in 10% formaldehyde (Fisher Scientific). Cells were washed with PBS and permeabilized with 0.01% Triton-X 100 (Sigma Aldrich) in PBS for 15 minutes at room temperature, followed by 2 washes in PBS-0.1% Tween 20 (Sigma Aldrich). Cells were then blocked in 1% bovine serum albumin (BSA) (Sigma Aldrich) for 1 hour at room temperature and washed with PBS-0.1% Tween 20. Cells were incubated with a primary antibody of anti-Human/Mouse/Rat Vinculin mouse monoclonal (R&D systems; MAB68961-SP) diluted to 1.25 μg/mL with 0.1% BSA in PBS-0.1% Tween 20 and a secondary antibody of 2 μL/mL Alexa Fluor 488-anti mouse in 0.1% BSA in PBS-0.1% Tween 20 (Abcam). Nuclei were stained with Hoescht 33342, trihydrochloride, trihydrate (Thermofisher; H3570) at 1 μg/mL for 30 minutes, and F-actin was stained with 1X Cytopainter 555 phalloidin (Abcam; ab176756) for 1 hour. Cells on coverslips were mounted onto microscope slides using Fluoromount Aqueous Mounting Media (Sigma Aldrich; F4680) and dried.

Fluorescence images of samples were obtained using a laser scanning confocal Olympus FV3000 microscope with 405 nm, 488 nm and 561 nm lasers. Images were processed using FiJi/ImageJ. The ‘Green Fire Blue’ pseudo-colour filter was applied for vinculin staining and ‘Red Hot’ for actin staining to observe the intensity and localization of biomolecule expression.

### Quantification of vinculin fluorescence

Using ImageJ image processing software, each cell membrane and nucleus were outlined by drawing boundaries around the area of interest. The mean fluorescence intensity per area within the outline was determined. The raw fluorescence intensity values of the nucleus were subtracted from the cytoplasmic values, as well as the subtraction of the nuclear area from the cytoplasmic area to obtain the ratio of nuclear to cytoplasmic fluorescence intensity and the area of the cytoplasm and nucleus, respectively. The fluorescence intensity was divided over the total area to obtain the fluorescence intensity/unit area for the nucleus and cytoplasm.

### PicoGreen cell adhesion study

Prior to the PicoGreen assay, all cells were aspirated of media, washed with PBS, and treated with 1X trypsin for 5 minutes at 37°C. The plates were shaken to detach cells and the suspended cells were aspirated off the wells. The PicoGreen assay (Quant-iT PicoGreen dsDNA reagent; Life Technologies) was then performed based on kit instructions to quantify the relative amount of cells left in each well based on fluorescence intensity.

MG-63 and HeLa cells were seeded into 24-well plates respectively at 35 000 cells/cm^2^ and grown to a confluency of 80% in 24 hours. MNP-PEI (0.35 μg Fe/μL) were added to cells different concentrations per well and placed on an oscillating magnet at 2 Hz and 0.2 mm amplitude for 30 minutes followed by a media change. Assay was performed after 24 hours of cell incubation.

For cells treated with MNPs, MG-63 cells were treated with oleic acid MNPs (0.35 μg Fe/μL) at different concentrations per well and placed on an oscillating magnet for 30 minutes after which one treatment group was assayed immediately whereas a media change was performed with 24-hour incubation on the other treatment group before the PicoGreen assay.

For cells treated with PEI polyplexes, MG-63 cells were treated with branched PEI 25kDa (1 mg/mL; Sigma) at different concentrations of PEI. The plates were placed on an oscillating magnet array for 30 minutes and one group was assayed subsequently, whereas the other plate was incubated for 24 hours before the assay.

### Quantitative real-time polymerase chain reaction (qPCR)

RNA extraction of MNP-PEI treated MG-63 cells was performed quantitative polymerase chain reaction (qPCR) analysis. The RNA extraction was performed according to the manufacturer’s instructions for the miRNeasy Mini Kit from Qiagen. Primers were designed via NCBI and supplied by Life Technologies.

Genes were quantified by one-step quantitative reverse transcription-polymerase chain reaction (q-PCR) using the QuantiFast SYBR Green RT-PCR kit (Qiagen, UK). A master mix was prepared by mixing 6.25 μl SYBR green, 1.25 μl of relevant primers (10 μM), and 0.125 μl enzyme per reaction tube, the master mix thoroughly mixed. 8.8 μl of master mix was loaded into individual wells containing 3.5 μL RNA sample (10 μM) and transferred to a Stratagene MX3005P real-time thermal cycler.

The thermal cycle consisted of 6 steps, including cDNA synthesis step (50°C for 10 min, 1 cycle), initial denaturation (95°C for 5 min, 1 cycle), denaturation (95°C for 10 sec, 40 cycles), primer annealing (60°C for 30 sec, 40 cycles), pre-programmed melt curve and fluorescent determination (95°C for 1 min, 60°C for 30 sec, 95°C for 30 sec, 1 cycle each). Replicate reactions for each target gene and housekeeping gene were performed for each sample. To quantify the expression difference among the samples, the ΔΔCt was calculated. The ΔCt value was calculated for each sample as the difference between the Ct for the target genes and the house keeping gene. ΔΔCt was measured as the difference between ΔCt values of an experimental sample and the control sample, the fold-change in gene expression was measured as 2^−(ΔΔCt)^.

qPCR products were separated and visualised on 2% agarose gels stained with 0.25μg/mL ethidium bromide (Sigma).

### Statistical analysis

Statistical analysis was conducted using OriginPro 2016. A 2-way ANOVA statistical analysis was performed for ferrozine assay data, vinculin fluorescence quantification data, and PicoGreen data. Statistical significance for qPCR was calculated by first performing an F-test on ΔCt values to determine equal or unequal variance. Based on the F-tests, the appropriate t-tests were performed on ΔCt values. Error bars are the calculated standard error of mean. The alpha value for statistical significance was set to 0.05. P values were denoted as *p<0.05, **p<0.005, ***p<0.0005 to indicate statistical significant differences between groups.

